# Gene Loss DB: A curated database for gene loss in vertebrate species

**DOI:** 10.1101/2025.05.26.656173

**Authors:** Gonçalo Espregueira Themudo, Raquel Ruivo, Raul Valente, Nádia Artilheiro, Diogo Oliveira, Inês Amorim, Bernardo Pinto, L. Filipe. C. Castro, Sergio Fernandes, Mónica Lopes-Marques

## Abstract

Molecular databases are essential resources for both experimental and computational biologists. The rapid increase in high-quality genome assemblies has led to a surge in publications describing secondary gene loss events associated with lineage-specific adaptations across diverse vertebrate groups. This growing volume of information underscores the urgent need for organized, searchable, and curated resources that facilitate data discovery, allow detection of broad evolutionary patterns, and support downstream analyses.

Currently, no existing database compiles manually curated and validated information on published secondary gene loss events. Here, we introduce the Gene Loss Database (GLossDB), a platform designed to centralize and present this data in an easy-to-search and user-friendly format (https://geneloss.org/).

GLossDB compiles gene loss events alongside supporting evidence, including the inferred mechanism of gene loss (exon deletion, gene deletion, loss of function mutation), the type of data used to support inactivation (genomic, transcriptomic, single/multiple individual sequence reads, synteny maps) and, when available, whether the event is shared across all lineages within a taxon. Each entry also includes a short excerpt from the original publication to provide context. This information is structured in the database to be searchable by species, gene, taxa, or by GO-terms linked to the gene in question.

The initial release of GLossDB focuses on cetaceans, a lineage with numerous gene loss events linked to aquatic adaptations. This first collection comprises 1866 gene loss events identified across 57 cetacean species. In addition, the database includes 1321 gene loss events from other taxa, which were also reported in the same studies and collected simultaneously.

## INTRODUCTION

The emergence of high-quality genomic data sheds light into the dynamic nature of genomes: shaped by the combination of selection, gene duplication, gene loss, and environmental adaptations. In vertebrates, genomic plasticity has been linked to phenotypic diversity, niche adaptation and trait innovation (1,2). Such growing wealth of information also uncovers critical knowledge gaps; particularly in the case of secondary gene loss, defined as the loss of ancestral functions in extant lineages (1,2). Gene loss is often poorly documented in genomic databases; yet, recent high-quality genome assemblies have highlighted its significance in evolutionary processes (2–4). Gene loss often occurs due to ORF disrupting mutations, deletions or even chromosomal rearrangements, with partial or total excision of gene sequences (3). Nonfunctional genes often share sequence similarity with functional homologs, which can lead to their misannotation as protein-coding by automated pipelines. This is more likely in species where supporting evidence, such as transcripts, proteins, or short reads from the same species, is limited, and thus genome annotation relies instead on data from closely related species (5). This persistent gap in genome annotation has fuelled numerous studies, notably in mammals with recently published genomes, focusing on the identification of secondary gene loss events often linked to lineage-specific adaptive traits (1–4,6–17).

Currently, a search on PubMed with the term “gene loss” yields thousands of results, most of which have been published since 2015, coinciding with the emergence of high-quality genomic data for numerous species. Such wealth of information brings out the timely need to organise and catalogue data to maximize searchability and streamline subsequent studies. Despite this, until now, there was no existing database collating curated and validated information regarding published secondary gene loss events. To find targeted information regarding gene loss episodes in a specific gene family or species, researchers must scavenge through numerous manuscripts addressing gene loss with varying degrees of evidence; and, more laboriously, through extensive supplementary material files, making it challenging to quickly and accurately compile data from multiple papers.

To address this issue, we present the Gene Loss Database (GLossDB), a user-friendly, curated, reliable, and efficient tool for navigating gene loss events. GLossDB does not focus on the annotation and/or identification of pseudogenes from non-curated sources, such as public genome databases (e.g. Ensembl and NCBI), setting itself apart from other existing tools such as PseudoChecker (18) and TOGA (19). Instead, this database collects data from published manuscripts previously subjected to peer-reviewed scrutiny and validation. Expert researchers (biocurators) extract data from these publications, which is then organized and deposited in the database. Most importantly, this database brings to the spotlight vast amounts of data often shadowed within the supplementary materials and thus with decreased visibility and searchability. This is particularly relevant for publications addressing large datasets e.g.(20,21). For example, one paper reports the inactivation of multiple vision-related genes in subterranean and other mammals adapted to low-light environments. While the main text highlights a few genes namely: *RBP3*, *OPN1SW/SWS1*, *GJ10*, *ARR3*, *CRB1*, *GRK7*, *GUCA1B*, and *GUCY2F* the study analysed 213 vision-associated genes, with the full list available in the supplementary material (20). Another example examines gene losses in the cetacean stem lineage where 11 genes are discussed in the main text, yet 74 gene losses are reported in the supplementary material (21).

To address these challenges, GLossDB aims to: (i) aggregate and organize gene loss data that is currently scattered across numerous publications, repositories and supplementary materials facilitating access to this information; (ii) promote a standardization of gene loss annotations (iii) promote data driven discovery by organizing curated data; and (vi) improve data traceability.

For the initial release of the database, we have selected the curation and aggregation of data from the Cetacea lineage. This choice was motivated by the recent availability of numerous cetacean genomes and the fact that this group exhibits a relatively high number of gene loss events, many of which have only been identified in recent years. In cetaceans, gene loss has been linked to key morphological and physiological traits associated with aquatic adaptation, e.g.,(13,16,21,22). Despite these changes, cetacean genomes retain a high level of sequence conservation with those of other mammals, which can lead to the frequent misannotation of pseudogenes as intact coding genes in public databases (5). Several studies have identified such pseudogenes and explored their potential adaptive relevance in cetaceans, including those associated with the loss of fur e.g., (12,21,23,24), the absence of sebaceous glands (7), alterations in skin structure, modifications in skin immunity, e.g.,(10,25–27), loss of tooth development in mysticeti species e.g., (16,17,28,29) loss of taste receptors, loss of visual receptors e.g., (13,30,31) among other cases. However, while many of these losses have been reported in the literature, the data remains scattered across numerous publications.

Here, we consolidate gene loss data for cetaceans from 55 published studies, providing a centralised, curated, searchable database resource - Gene Loss Database (GLossDB).

## METHODS

### Database implementation

The Gene Loss Database frontend data exploration interface and the backend data curation interface for curators were specifically developed for this purpose. GLossDB was built using Laravel, a robust and flexible PHP framework for web application development. The data storage engine employed was MySQL, a widely used and free relational database management system. For the backend administration interface, AdminLTE, a popular open-source admin dashboard template, was used to provide a responsive and customizable UI. To enhance the usability of the web interface, Select2 (available at http://select2.org), a jQuery-based replacement for select boxes, and DataTables, a plugin for enhancing HTML tables, were integrated. For the frontend service, Bootstrap, a popular CSS framework for responsive and mobile-first web development, was utilized. Additionally, data visualization capabilities were implemented using Chart.js, an open-source JavaScript library for creating flexible and interactive charts.

GLossDB also integrates third-party APIs from NCBI, PantherDB, and GO-terms (32–34) to ensure data consistency, enrich data retrieval and analysis capabilities.

### Data collection, curation and quality control

Published manuscripts on gene loss were collected through a comprehensive search in PubMed and PubTator3 (35), excluding reviews. The search was conducted using a combination of the following terms: “gene loss”, “pseudogene”, “gene inactivation”, “gene disruption”, “gene deletion”, “cetacea” and “marine mammal,” along with the Boolean operator AND. The retrieved manuscripts were compiled into a non-redundant list, which was then manually reviewed to identify studies describing gene loss events in cetaceans. Manuscripts meeting the inclusion criteria underwent full-text review and were incorporated into the database as annotation jobs. Additionally, curators could suggest relevant manuscripts that were not identified in the initial search, which were also included.

Gene loss data curation was performed by expert curators who read the selected manuscripts and extracted gene loss information in three steps. In the first step, all structured data is collected by completing the Gene loss (GLoss) annotation forms, either by providing NCBI identifiers or by selecting the correct option from the provided list (Table1).

**Table: 1.** Structured information requested in a GLoss annotation.

| Information requested | Curator reply |
| --- | --- |
| Identification of the gene reported to be lost | **Reply:** NCBI Gene ID of the reference gene |
| Identification of the species in which the gene is reported to be lost | **Reply:** NCBI Tax ID of the species in which the gene is lost |
| Determination of the type of gene loss | **Options:** Full, Polymorphic, Undetermined |
| Identification of the gene loss mechanism | **Options:** Gene deletion, exon(s) deletion, Loss of Function-LOF (frameshift, premature stop, abolishment of canonical splice sites), regulatory region mutation, other. |
| Identification of the presented evidence to validate gene loss | **Options:** Multiple individual SRA, Single Individual SRA, Genomic, Transcriptomic, Genomic & Synteny maps, Genomic & Transcriptomic, PCR & Sanger Sequencing Multiple, PCR & Sanger Sequencing Single, Sequence Trace Archive |
| Indicate if the gene loss is shared in all organisms of a specific taxon lineage specific | **Options:** Yes; No |
| Accession number of the lost gene if available | **Reply:** Accession number (optional) |

In step two, curators select short excerpts from the manuscript that provide additional context for the gene loss annotations. These excerpts (statements) are then incorporated into the gene loss annotation and categorized based on the type of information they contain. The categories include “Mutational Description”, “Functional”, “Phenotypic”, “Timing of Loss”, “Methodology & Validation” and “Other”. In step three, curators can provide critical insights into a specific gene loss event that may not be immediately evident from the manuscript. These observations are recorded in the “Curator Observations” section as unstructured free-text data.

After collecting all data from a specific annotation job, curators submit the full annotation job containing multiple GLoss annotations for quality control. Quality control consists of two rounds of data validation. The first round is performed programmatically/computationally, ensuring that all required fields are completed and that stable identifiers such as gene ID and tax ID are used appropriately, preventing duplicate annotations within the same job. The second round involves manual validation by trained database curators, who review each annotation job to ensure consistency and completeness (Figure 1). Only after successfully undergoing both validation steps is the annotation job completed and then published in the database.

**Figure 1:**
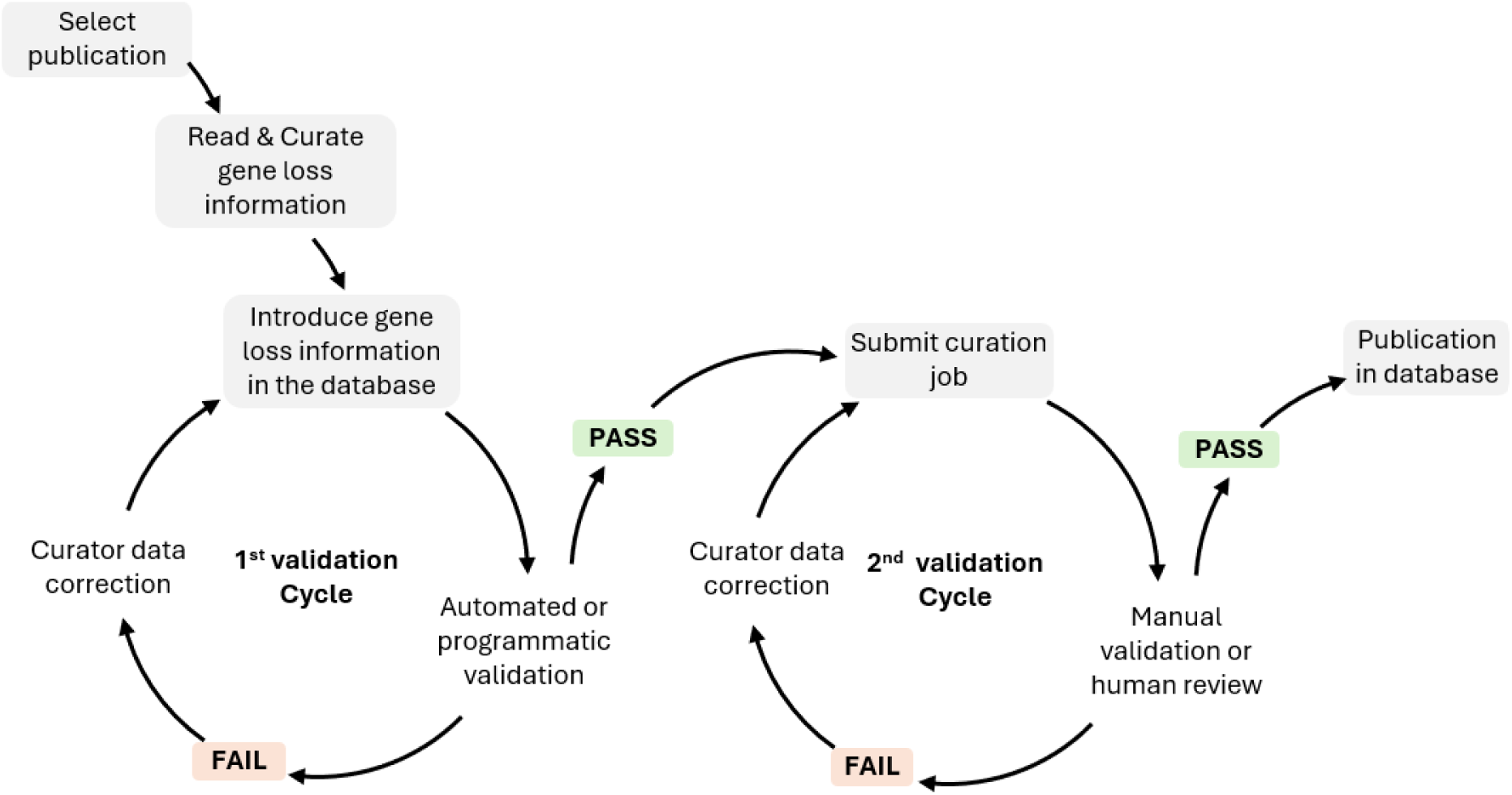
Database curation workflow and validation steps.

### Data structure and searchability

The fundamental unit of the Gene Loss Database is a Gene Loss Annotation or GLoss Annotation, which documents the loss of a single gene in a specific species. Each GLoss Annotation is linked to a reference manuscript and to a Reference Gene, the highest-ranking unit in the database’s organizational structure. A Reference Gene consolidates all GLoss Annotations related to that gene, regardless of the species or manuscript in which the loss was reported (Figure 2A).

**Figure 2:**
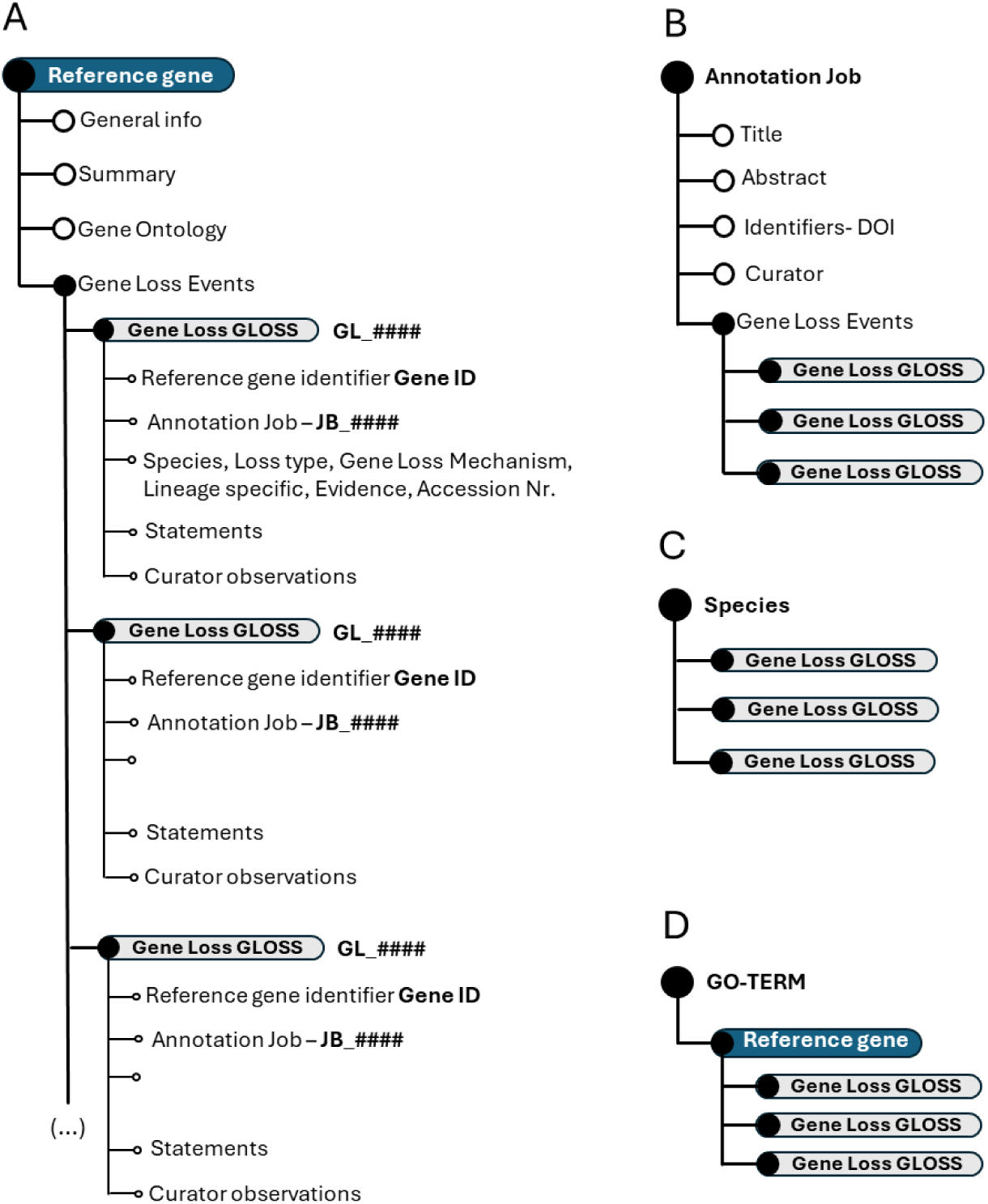
Schematic representation of the Gene Loss database structure. (A) Hierarchical organization of the overall database (B) Data structure when explored by annotation job (C) Data structure when explored by species (D) Data organization when focused on GO-terms.

In most cases, the Reference Gene corresponds to the coding ortholog in the human genome. However, if the human gene is non-coding, a coding orthologue from another model species may be selected as the reference.

The reference gene is selected using a stable identifier -Gene ID (structured information - Table 1), allowing the automatic retrieval via API from the NCBI gene database of general information linked to the gene including: gene summary, symbol, aliases, and GO-terms, as well as paralogues from Panther Knowledgebase (32).

The Gene Loss Database has implemented a dynamic data structure, cross-linking, and use of unique identifiers to enhance data searchability and user-friendliness. As a result, the data structure may vary depending on the user’s starting point. Yet, regardless of the user starting point, each GLoss annotation retains all associated identifiers, ensuring comprehensive traceability. Each GLoss Annotation includes the following automatically linked identifiers upon creation:

i. Reference Gene Identifier (Gene ID): Identifies the reference gene associated with the annotation
ii. GLoss Identifier (GL_######): A unique, six-character alphanumeric identifier preceded by “GL”, generated automatically by the database;
iii. Annotation Job Identifier (JB_######): A unique, six-character alphanumeric identifier preceded by “JB” linking each GLoss annotation to a reference publication.

Exploring the Gene Loss Database by Annotation Job reveals all GLoss annotations associated with a single annotation job (Figure 2B). When exploring gene loss data by Species, all GLoss annotations linked to a specific species are displayed, regardless of their annotation job of origin (Figure 2C). Finally, organizing the data by GO term presents all GLoss annotations associated with a specific GO term, which is linked to a corresponding term in the reference gene (Figure 2D).

## RESULTS

### Database Usage – browsing and targeted search

The Gene Loss Database can be explored either by browsing all available data or through targeted searches. Users can browse the data through icons on the homepage (Figure 3A). Browsing by selecting the pseudogene icon will return a list of all GLoss annotations in the database. Browsing by selecting the species icon provides a list of all species with at least one GLoss annotation, allowing users to select a species and view all associated GLoss annotations. Browsing by publication displays a list of curated publications in the current version of the database, showing the total number of GLoss annotations extracted from each publication along with direct links to the published article (Supplementary Table 1).

**Figure 3.**
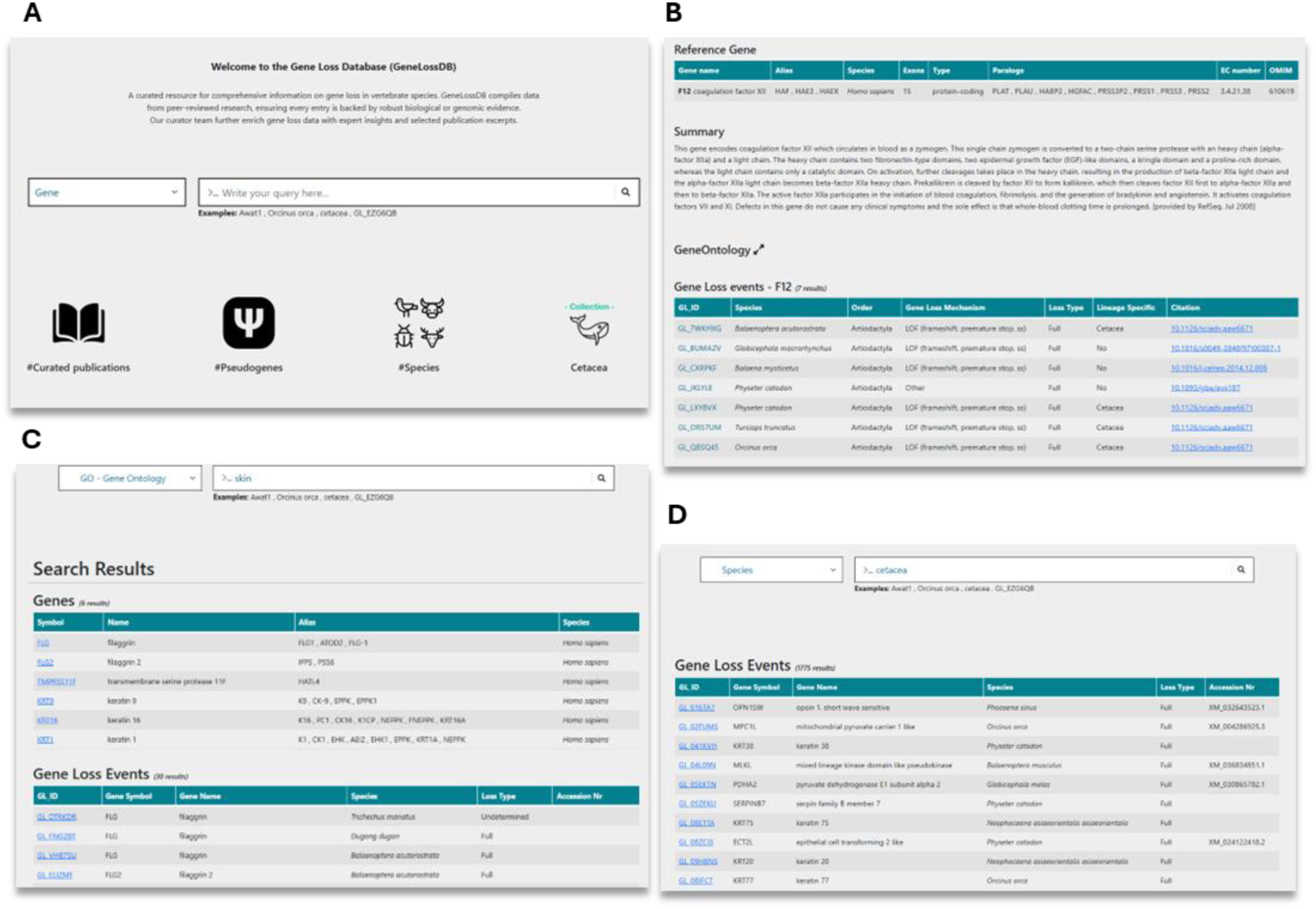
Overview of the Gene Loss Database and search methods. **(A)** Homepage with navigation options. **(B)** Reference gene page displaying GLoss annotations linked to a specific gene. **(C)** Search results for a Gene Ontology (GO) term query, showing associated genes. **(D)** Search results for a species query, listing gene loss events reported for the selected species.

Users may also perform targeted searches using the search box. To begin a search, they must first select a category from the drop-down menu, which includes Gene, Species, GLossID, and GO-terms (Figure 3A). To enhance user experience and facilitate data exploration, the Gene Loss Database supports “partial exact matching”, meaning users do not need to enter the full search term to obtain results, but correct spelling is required. When searching by gene, users can enter a gene symbol, gene alias, full gene name, or partial gene name. This returns a list of reference genes, and upon selection, users are redirected to a page containing reference gene details and all linked GLoss annotations (Figure 3B). Searching by species allows input of a species name, partial species name, common name, order, or infraorder, retrieving GLoss annotations associated with species matching the search terms. Users can also explore gene loss data from a functional perspective using the GO term search, where keywords should correspond to full or partial GO terms or GO term ID numbers (Figure 3C and 3D). This returns a list of reference genes linked to the specified GO term, and selecting a gene provides access to its associated GLoss annotations. Finally, searches can also be conducted using GLoss identifiers. By selecting the GLossID search option and entering a specific GLossID, users retrieve the corresponding GLoss annotation directly.

Each GLoss annotation page is structured into several sections to ensure clarity and ease of navigation. At the top, the general information section (Figure 4A) provides a header displaying the gene symbol and the species in which the gene is reported as lost. Below this, additional details such as cross-links to the reference gene and the corresponding annotation job are included (Figure 4B). The next sections focus on describing the gene loss event, these combine structured data (Figure 4C), including the GLossID, species, gene loss mechanism, loss type, supporting evidence, and lineage specificity, with semi-structured data (Figure 4D) in the form of text excerpts selected by the curator and extracted from the curated manuscript. These excerpts may be classified into 6 types depending on the type of information included here: (i) Phenotypic – excerpts addressing the phenotypic outcome associated to the reported gene loss, (ii) Functional - excerpts describing the function of the gene and corresponding protein encoded (iii) Timing of Loss-excerpts indicating the approximate timing of gene loss (iv) Mutation description – excerpts with general description of the identified ORF disruption mutations, (v) Methodology & validation – excerpts with a general description of the methods used to identify and validate the reported gene loss event and (vi) Other-text segments selected from the manuscript that the curator deemed as essential to provide context to the GLoss annotation and that cannot be classified in the previous types. Following this, the curator observations section (Figure 4E) contains unstructured data, where expert curators offer specific insights and additional context regarding the gene loss event. This section highlights any critical details that may not be explicitly stated in the manuscript but are relevant for data interpretation. Finally at the bottom of the page, the related GLosses section (Figure 4F) lists other instances in which the same gene was reported as lost in the same species but in different curated manuscripts. This feature helps users identify repeated findings across independent studies, further supporting the reliability of gene loss reports. This also ensures complete coverage, as all gene losses reported in each publication are curated, which may result in multiple independent annotations for the same gene and species. While this introduces a degree of redundancy, it also increases the robustness of the database by enabling independent corroboration of findings across diverse sources.

**Figure 4:**
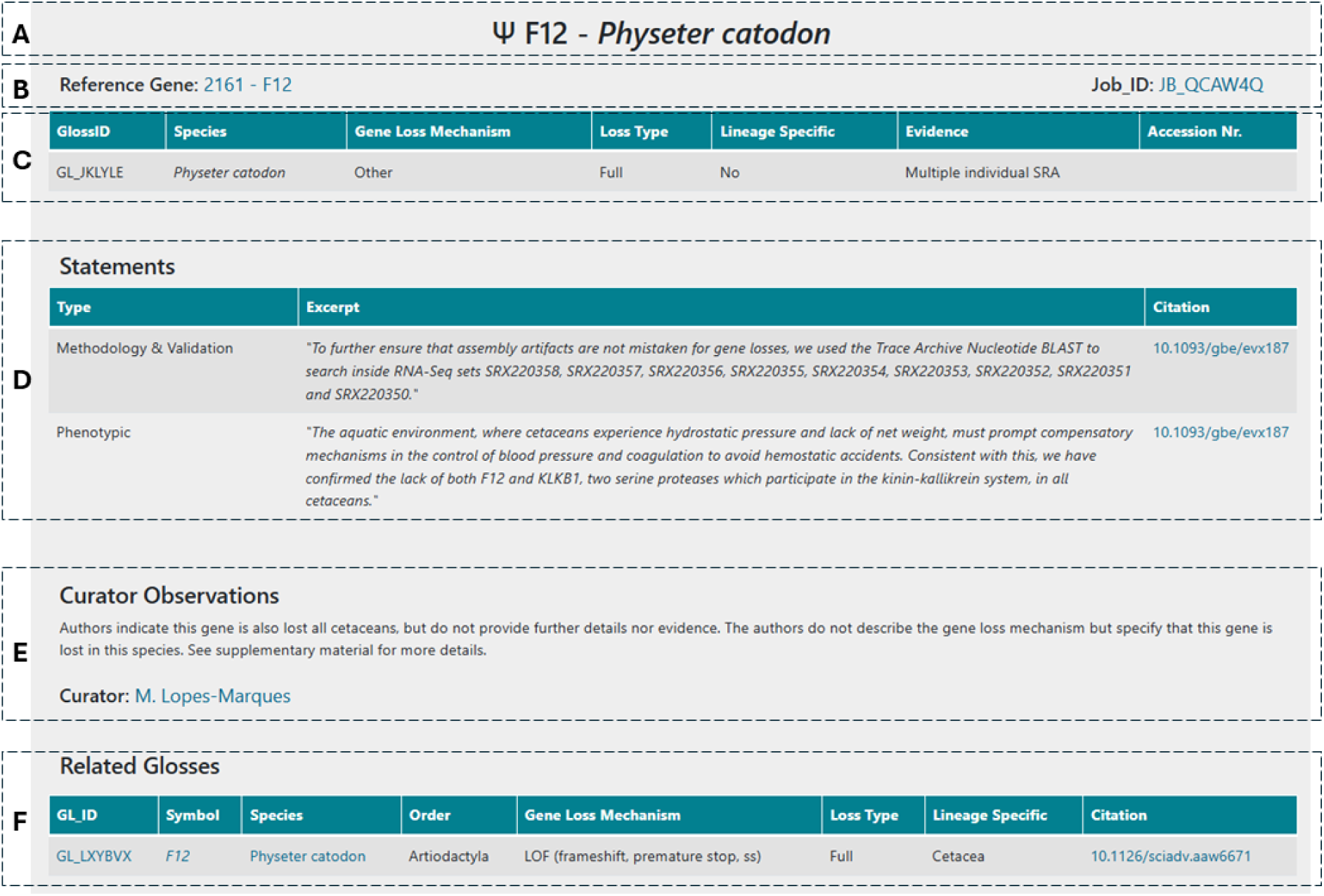
Example of the general structure of a GLoss annotation page.

### Database Contents

Although this collection focuses on gene loss in cetacean species, all gene loss events reported in the selected publications were annotated, leading to a spillover into other mammalian orders. Currently, the Gene Loss Database contains curated data from 55 publications analysed within the scope of the cetacean collection (see Supplementary File 1). This curation effort resulted in 3187 gene loss annotations across 443 genes in 359 mammalian species (including subspecies), with representatives from 22 mammalian orders (Table 2). Not surprisingly, the mammalian order with the highest number of species and GLoss annotations is Artiodactyla, which includes the cetacean infraorder. Following Artiodactyla, we find Carnivora with 209 GLoss annotations from 80 species, and Chiroptera with 152 GLoss annotations in 39 species (Table 2). It is important to note that insofar the only nearly complete collection in the database pertains to cetaceans; yet, GLoss annotations are expected to grow to cover the complete mammalian catalogue of gene loss events.

**Table 2:** Overview of the curated gene loss data included. * Not included in NCBI taxonomy but a recognised order.

| Order | N. of species | N. of GLoss annotations |
| --- | --- | --- |
| **Artiodactyla** | 109 (including 57 cetacea) | 2058 (including 1866 cetacea) |
| **Afrosoricida\*** | 2 (*Chrysochloris asiatica*, *Echinops telfairi*) | 49, 17 |
| **Rodentia** | 23 | 150 |
| **Carnivora** | 82 | 209 |
| **Primates** | 26 | 113 |
| **Perissodactyla** | 14 | 38 |
| **Pholidota** | 4 | 98 |
| **Dermoptera** | 1 (*Galeopterus variegatus*) | 8 |
| **Pilosa** | 8 | 27 |
| **Eulipotyphla** | 7 | 73 |
| **Chiroptera** | 40 | 152 |
| **Scandentia** | 1 (*Tupaia chinensis*) | 10 |
| **Proboscidea** | 6 | 34 |
| **Sirenia** | 4 | 63 |
| **Cingulata** | 5 | 41 |
| **Tubulidentata** | 1 (*Orycteropus afer*) | 14 |
| **Lagomorpha** | 10 | 22 |
| **Hyracoidea** | 2 | 5 |
| **Macroscelidea** | 1 (*Elephantulus edwardii*) | 1 |
| **Monotremata** | 2 | 2 |
| **Dasyuromorphia** | 1 (*Sarcophilus harrisii*) | 1 |
| **Diprotodontia** | 1 (*Vombatus ursinus*) | 1 |

Concerning gene loss mechanisms and evidence, data curation revealed that the most frequent gene loss mechanism reported in over 53% of the GLoss annotations was loss of function (LOF) mutations. These include frameshift mutations, mutations that alter canonical donor and acceptor splicing sites, and premature stop codons. This was followed by gene loss mechanisms included in the ‘other’ category (circa 19.5%), which include mutations that abolish the start codon, as well as cases in which the exact mechanism of gene loss was not specified by the authors of the original paper. Also, gene and/or exon(s) deletions were evident among the gene loss mechanisms mentioned by the authors (circa 16.4%, circa 10.3%, respectively) (Figure 5). In cases where gene loss is due to multiple mechanisms, such as the combination of LOF mutations and exon deletions, the Gene Loss Database prioritizes gene loss mechanisms that are shared across multiple species. For example, a shared premature stop codon will be given preference over an exon deletion observed in only one species within the same group. If no shared mutations are identified across several species under analysis, preference will be given to the most frequently observed mutation, followed by the first reported mutation that appears in the 5’region of the canonical isoform of the gene. It is important to note that while one gene loss mechanism is annotated in the database, the gene in question may carry other mutations and forms of gene erosion, as is expected in pseudogenes e.g., (7,11,36).

**Figure 5.**
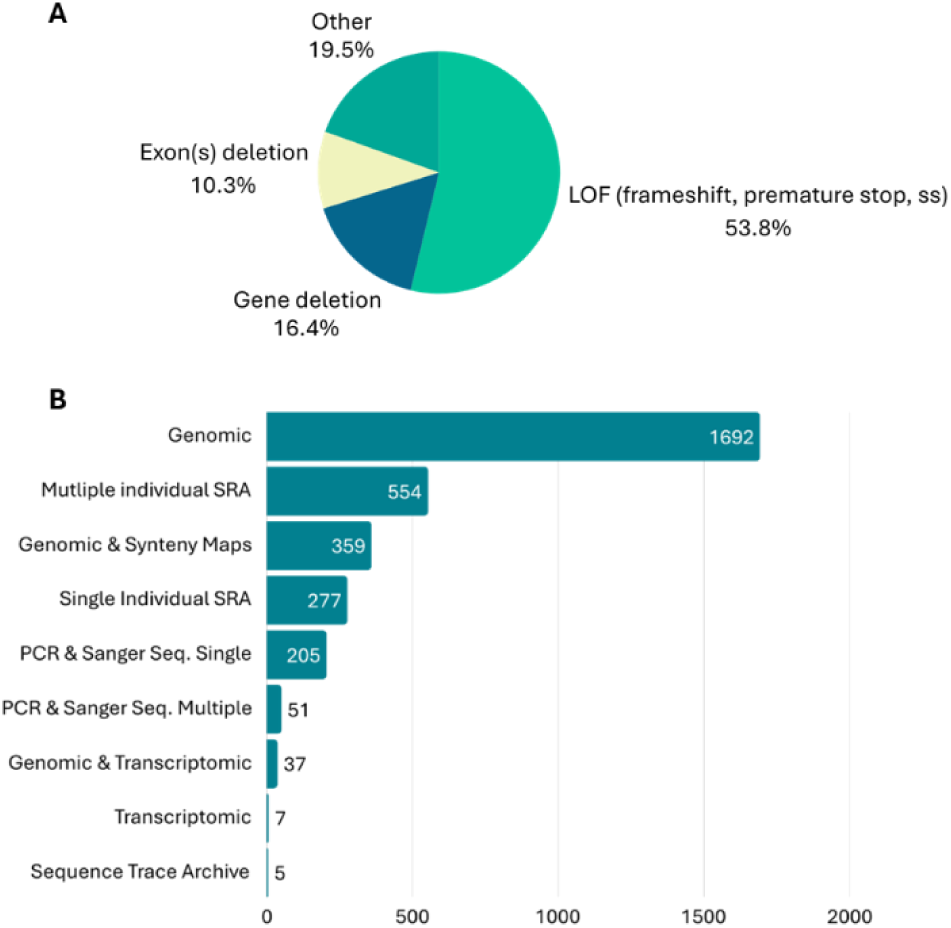
**A-**Mutational spectrum of main gene loss mechanisms reported in the current release of Gene Loss Database **B**- Gene loss evidence reported in the current release Gene Loss Database.

When considering the evidence provided by the authors to support claims of gene loss, genomic evidence was the most frequently mentioned. This refers to the identification of gene loss mechanisms using publicly available genomic data or genome assembly of the species in question. Often, authors combined genomic evidence with synteny maps and transcriptomic data to further support their findings. In addition to genomic evidence, many authors also provided multiple species SRA (Sequence Read Archive) datasets or single-species SRA datasets. Finally, in some cases, authors provided PCR and Sanger sequencing data from independent samples to validate the identified mutations (Figure 5). In cases where the authors present multiple forms of evidence supporting the identification of the gene loss mechanism, curators select the strongest form of evidence for inclusion in the database. This typically includes multi-species SRA datasets or other evidence validating the existence of the same mutation in multiple independent samples from the same species.

Gene loss was classified into three main categories: Full, Polymorphic, and Undetermined. A gene loss event was classified as “Full” when the gene in question was lost in all individuals of a specific species, indicating that the ORF-disrupting mutation has reached full fixation in that species. To validate this, curators screened the manuscripts for at least one of the following pieces of evidence: (i) the gene in question presents multiple LOF mutations, or (ii) if a single mutation is reported, the authors did not find evidence that this variation may be polymorphic, and/or (iii) the identified mutations were conserved with those observed in a sister species. A gene loss event was classified as “Polymorphic” when a single ORF-disrupting mutation was present in some individuals of a species but absent in others, indicating that the mutation had not reached full fixation (36,37). To confirm this classification, curators validated whether (i) the mutation was observed in a subset of analysed individuals from the same species and/or (ii) the authors explicitly stated that the gene in question was a polymorphic pseudogene in the target species. Some examples of polymorphic pseudogenes included in the current release of the database are *OPN1SW* in *Delphinapterus leucas* and *Phocoenoides dalli* (22), *MMP20* in *Kogia breviceps* (16), and *CNGA3* in *Eubaleana glacialis* (30). Finally, gene loss was classified as “Undetermined” when the authors explicitly expressed uncertainty about the gene’s coding status and when the manuscript lacked sufficient evidence to fully support the claim of gene loss. Examples include cases where the ORF-disrupting mutation was located in the last exon or near the end of the gene and truncating mutations are present but do not rule out protein functionality, as was the case for *TCHHL1* and *FLG2* in *Dugong dugong* and *Trichechus manatus* (38) and *IL20* in *Trichechus manatus* (10), or the authors did not identify any ORF-disrupting mutations but instead found multiple missense mutations affecting critical residues as in the case of *CORT* in *Pontoporia blainvillei* (39). In the current database collection, a total of 3138 gene loss annotations were classified as full gene loss events, 16 were reported as polymorphic, and 33 were classified as undetermined.

When analysing the genes reported as lost, an overall review of the database shows that *PCSK9* was the gene most frequently reported as lost, with 186 GLoss annotations emerging from two independent large multispecies studies (4,15) and with a single GLoss annotation in a third study (4). Since these annotations come from independent sources, redundant gene loss reports for *PCSK9* in the same species were detected in 14 species. These duplicate annotations are referred to as “*related Glosses”* in the database (see Figure 4). Following *PCSK9,* the genes with the highest number of gene loss annotations are *MTNR1B*, *CORT,* and *UCP1,* with 67, 63, and 53 GLoss annotations, respectively.

### The cetacean collection

Currently, the cetacean dataset comprises 1866 GLoss annotations referencing the loss of 314 genes in 57 cetacean species, including 15 Mysticeti and 42 Odontoceti species (Figure 6A). The species with the highest number of GLoss annotations are *Tursiops truncatus*, *Balaenoptera acutorostrata*, *Physeter macrocephalus*, and *Orcinus orca* each with over 200 annotations. It is important to note that this high number of reported gene losses in these species primarily reflects the early availability of their genome assemblies in public databases and the extensive research conducted on these organisms, rather than a greater propensity for gene loss. Furthermore, this does not imply that these genes remain intact in other cetaceans, which have not been investigated.

**Figure 6:**
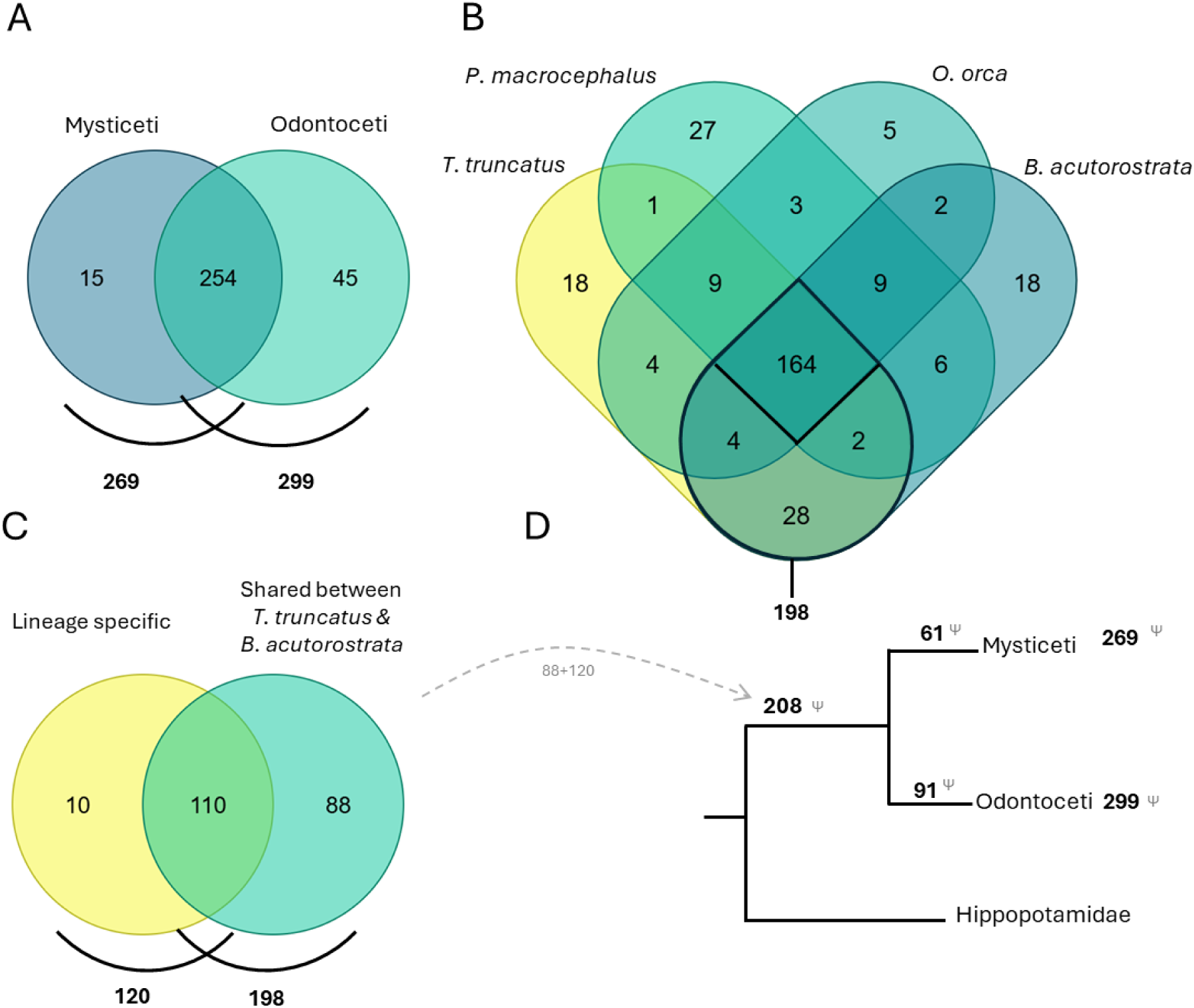
**A** – Comparison of genes lost in all Mysticeti and Odontoceti species annotated in the GLossDB**. B -** Comparison of genes lost in 4 species with the highest number of Gloss annotations. **C -** Comparison of genes lost shared between *T. truncatus* and *B. acutorostratata* (198) with genes reported in the literature to be lost in the cetacean ancestor, lineage-specific (120). **D -** Schematic representation of gene loss data in cetacean lineage. (Venn diagrams prepared using dataset intersections at molbiotools.com)

A comparative analysis of gene losses across all four species identified 164 shared lost genes. When focusing on the two species with the highest number of GLoss annotations, *T. truncatus* (Odontoceti) and *B. acutorostrata* (Mysticeti), each representing a major cetacean lineage, the number of shared gene losses increases to 198 (Figure 6B). The occurrence of these gene losses in both lineages suggests that they may have taken place before the divergence of Odontoceti and Mysticeti or depict parallel adaptation scenarios under similar environmental constraints. In the current cetacean data collection, curators aimed to determine whether reference manuscripts reported gene losses in the cetacean ancestor and/or if these losses could be attributed to a single mutational event (lineage-specific) in the cetacean ancestor. Through this analysis, a total of 120 genes were identified as having been lost in the ancestral cetacean lineage. When comparing this list with the 198 genes found to be lost in both *T. truncatus* and *B. acutorostrata*, we identified an overlap of 110 genes. The observed overlap supports the hypothesis that genes absent in both Mysticeti and Odontoceti were predominantly lost in their common ancestor, though independent early losses in each lineage cannot be excluded in all cases. Based on this, we infer that an additional 88 genes, previously not classified as lineage-specific losses are also potentially lost in the cetacean ancestor (Figure 6C). This brings the total number of putatively lost genes in the cetacean lineage to 208 genes.

To gain insight into the main biological processes affected by the loss of these genes, a gene ontology (GO) term analysis was performed (32,33). For this analysis, the 208 genes reported as pseudogenized in cetaceans in GLossDB were compiled and queried against gene ontology databases to test for overrepresentation (32,33). The results revealed significant enrichment in several biological processes (Table 3). As expected, many of these processes are linked to specific adaptations of these mammals to the aquatic environment. For example, these include the remodeling of the melatonin biosynthetic and metabolic process, which has been linked to the altered circadian rhythm of cetaceans (8), the remodeling of skin phenotype, characterised by the loss of fur and sebaceous glands, and modifications in the skin barrier (7,12,23,24). We also observed a significant enrichment of genes associated with keratinocyte and epidermal development, as well as cell differentiation. These findings support the hypothesis of extensive evolutionary modifications in skin development, which likely occurred in ancestral lineages, consistent with previous reports (4,7,10,12). Additionally, we observed a significant enrichment of genes related to sensory perception, such as taste, possibly reflecting adaptations to aquatic diets, or the visual sensory system, suggesting adaptations to underwater vision (20,30).

**Table 3:**
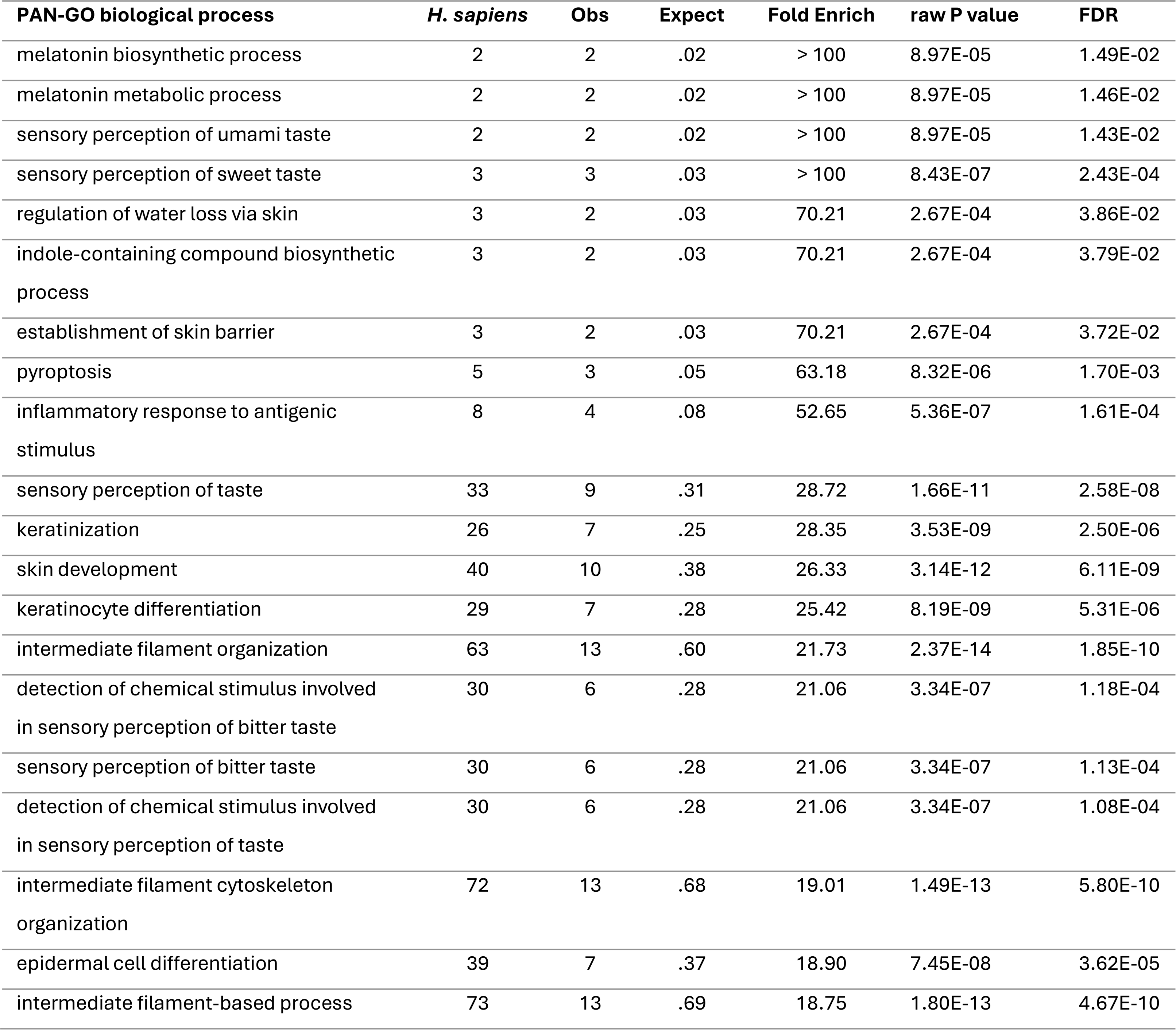
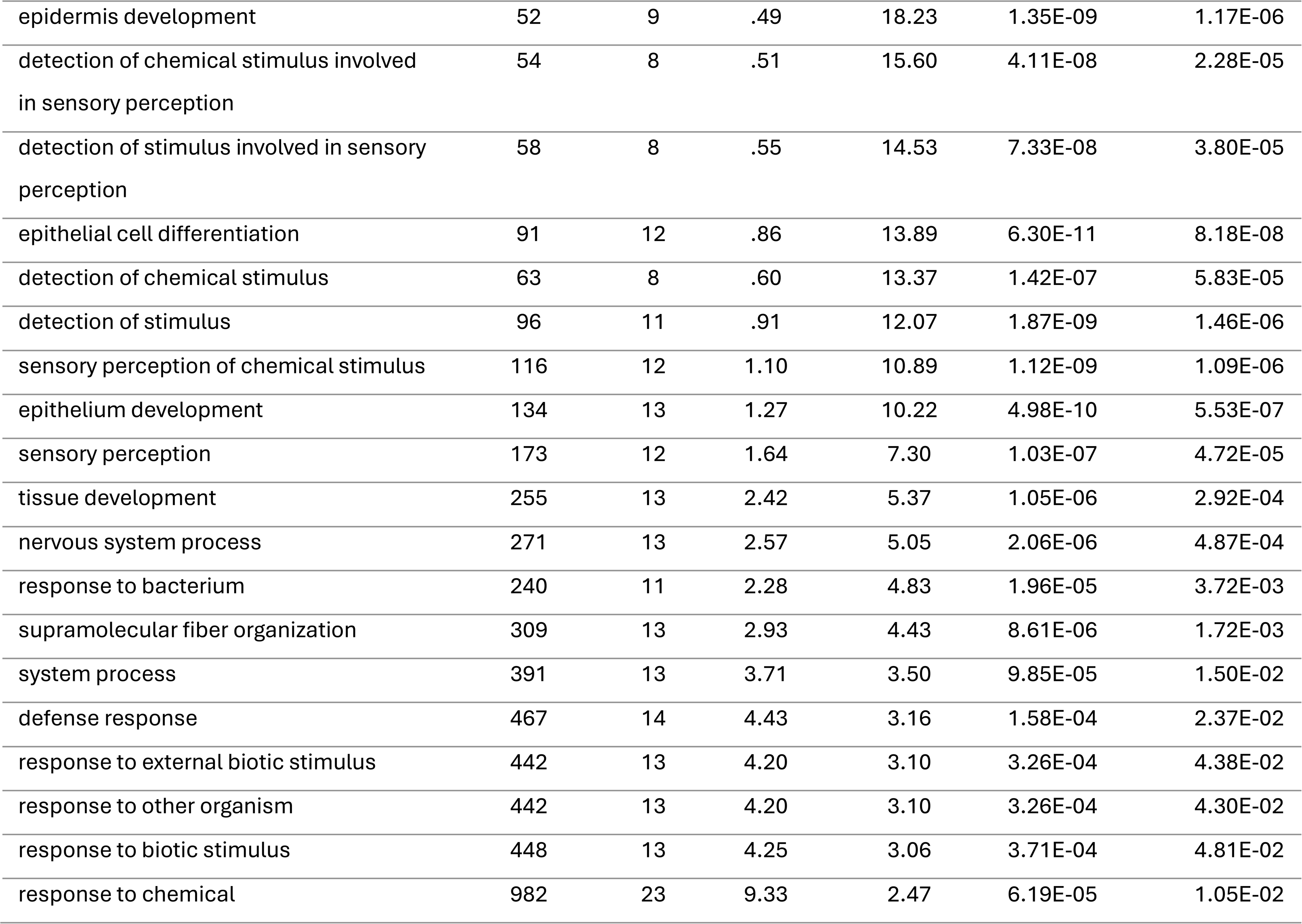
GO Term Biological process enrichment analysis.

| PAN-GO biological process | *H. sapiens* | Obs | Expect | Fold Enrich | raw P value | FDR |
| --- | --- | --- | --- | --- | --- | --- |
| melatonin biosynthetic process | 2 | 2 | .02 | > 100 | 8.97E-05 | 1.49E-02 |
| melatonin metabolic process | 2 | 2 | .02 | > 100 | 8.97E-05 | 1.46E-02 |
| sensory perception of umami taste | 2 | 2 | .02 | > 100 | 8.97E-05 | 1.43E-02 |
| sensory perception of sweet taste | 3 | 3 | .03 | > 100 | 8.43E-07 | 2.43E-04 |
| regulation of water loss via skin | 3 | 2 | .03 | 70.21 | 2.67E-04 | 3.86E-02 |
| indole-containing compound biosynthetic process | 3 | 2 | .03 | 70.21 | 2.67E-04 | 3.79E-02 |
| establishment of skin barrier | 3 | 2 | .03 | 70.21 | 2.67E-04 | 3.72E-02 |
| pyroptosis | 5 | 3 | .05 | 63.18 | 8.32E-06 | 1.70E-03 |
| inflammatory response to antigenic stimulus | 8 | 4 | .08 | 52.65 | 5.36E-07 | 1.61E-04 |
| sensory perception of taste | 33 | 9 | .31 | 28.72 | 1.66E-11 | 2.58E-08 |
| keratinization | 26 | 7 | .25 | 28.35 | 3.53E-09 | 2.50E-06 |
| skin development | 40 | 10 | .38 | 26.33 | 3.14E-12 | 6.11E-09 |
| keratinocyte differentiation | 29 | 7 | .28 | 25.42 | 8.19E-09 | 5.31E-06 |
| intermediate filament organization | 63 | 13 | .60 | 21.73 | 2.37E-14 | 1.85E-10 |
| detection of chemical stimulus involved in sensory perception of bitter taste | 30 | 6 | .28 | 21.06 | 3.34E-07 | 1.18E-04 |
| sensory perception of bitter taste | 30 | 6 | .28 | 21.06 | 3.34E-07 | 1.13E-04 |
| detection of chemical stimulus involved in sensory perception of taste | 30 | 6 | .28 | 21.06 | 3.34E-07 | 1.08E-04 |
| intermediate filament cytoskeleton organization | 72 | 13 | .68 | 19.01 | 1.49E-13 | 5.80E-10 |
| epidermal cell differentiation | 39 | 7 | .37 | 18.90 | 7.45E-08 | 3.62E-05 |
| intermediate filament-based process | 73 | 13 | .69 | 18.75 | 1.80E-13 | 4.67E-10 |
| epidermis development | 52 | 9 | .49 | 18.23 | 1.35E-09 | 1.17E-06 |
| detection of chemical stimulus involved<br>in sensory perception | 54 | 8 | .51 | 15.60 | 4.11E-08 | 2.28E-05 |
| detection of stimulus involved in sensory<br>perception | 58 | 8 | .55 | 14.53 | 7.33E-08 | 3.80E-05 |
| epithelial cell differentiation | 91 | 12 | .86 | 13.89 | 6.30E-11 | 8.18E-08 |
| detection of chemical stimulus | 63 | 8 | .60 | 13.37 | 1.42E-07 | 5.83E-05 |
| detection of stimulus | 96 | 11 | .91 | 12.07 | 1.87E-09 | 1.46E-06 |
| sensory perception of chemical stimulus | 116 | 12 | 1.10 | 10.89 | 1.12E-09 | 1.09E-06 |
| epithelium development | 134 | 13 | 1.27 | 10.22 | 4.98E-10 | 5.53E-07 |
| sensory perception | 173 | 12 | 1.64 | 7.30 | 1.03E-07 | 4.72E-05 |
| tissue development | 255 | 13 | 2.42 | 5.37 | 1.05E-06 | 2.92E-04 |
| nervous system process | 271 | 13 | 2.57 | 5.05 | 2.06E-06 | 4.87E-04 |
| response to bacterium | 240 | 11 | 2.28 | 4.83 | 1.96E-05 | 3.72E-03 |
| supramolecular fiber organization | 309 | 13 | 2.93 | 4.43 | 8.61E-06 | 1.72E-03 |
| system process | 391 | 13 | 3.71 | 3.50 | 9.85E-05 | 1.50E-02 |
| defense response | 467 | 14 | 4.43 | 3.16 | 1.58E-04 | 2.37E-02 |
| response to external biotic stimulus | 442 | 13 | 4.20 | 3.10 | 3.26E-04 | 4.38E-02 |
| response to other organism | 442 | 13 | 4.20 | 3.10 | 3.26E-04 | 4.30E-02 |
| response to biotic stimulus | 448 | 13 | 4.25 | 3.06 | 3.71E-04 | 4.81E-02 |
| response to chemical | 982 | 23 | 9.33 | 2.47 | 6.19E-05 | 1.05E-02 |

## CONCLUSIONS

One of the major challenges in the genomic era is the ability to compile and analyse increasingly large volumes of data. The rapid growth in available genomic sequences has expanded the raw material for identifying evolutionary and functional patterns. At the same time, however, it has exposed important analytical limitations, particularly the frequent misannotation of secondary gene loss by automated annotation pipelines (5). As a result, gene loss research is gaining momentum not only for its evolutionary relevance but also, because it offers valuable insights into biological processes by acting as a source of natural knockouts. Although efforts have been made to streamline the annotation and/or identification of pseudogenes (18,19), secondary gene loss events have been largely reported and validated in scattered manuscripts, with no centralised database systematically collecting and integrating this information.

The Gene Loss Database addresses this gap by systematically aggregating and organizing gene loss information in a single and user-friendly resource making it compatible with FAIR principles and Open Science (40). By centralizing gene loss research, we overcome the archival fragmentation of data: facilitating comparative analyses, improving data discoverability, traceability and enabling novel connections that might otherwise be overlooked.

While the current collection focuses on cetaceans, the database is expanding, to incorporate additional taxonomic groups with relevance to evolutionary and health research. Future collections will include species that serve as natural models for human disease, further bridging the gap between evolutionary genetics and biomedical applications. As more data becomes available, this resource will provide deeper insights into gene function, natural knockouts, and disease mechanisms, reinforcing the importance of gene loss studies in both evolutionary and biomedical sciences.

## OPEN INVITATION

With the publication of this database, we extend an open invitation to the research community to contribute. Researchers are encouraged to identify and communicate relevant articles for curation and inclusion in the Gene Loss Database via our message box on https://geneloss.org/about. Alternatively, researchers can contact us to propose and lead a curation effort for a specific species or group of species within their research focus or interest.

## DATA AVAILABILITY

GLossDB does not require user registration and is available online at https://geneloss.org.

## SUPPLEMENTARY DATA

Supplementary Table 1 – Curated publications.

Supplementary Table 2 - Number of gloss annotations in Cetacea by species.

## AUTHOR CONTRIBUTIONS

**Monica Lopes-Marques:** Conceptualization, Formal analysis, Methodology, Validation, Writing—original draft, preparing final manuscript and submission. **Sergio Fernandes:** Software development and implementation of the full database, including back-end and front-end office. Server management and maintaining the database online.

**Mónica Lopes-Marques, Gonçalo E. Themudo, Raul Valente, Nadia Artilheiro, Inês Amorim, Diogo Oliveira, Bernardo Pinto:** Expert curators for data collection and input into the database, review and validation of data, and critical contributions to the manuscript.

**Gonçalo E. Themudo, Raquel Ruivo, and Filipe Castro:** Critical feedback and optimization of the database workflow, writing and review of the manuscript. All authors read and reviewed the final version of the manuscript.

## ACKNOWLEDGEMENTS

We acknowledge the support of CORAL AI, an instrumental tool that fast-tracked the reading and extraction of data from numerous manuscripts. We also acknowledge NCBI, PantherDB, and Gene Ontology for their API interfaces, which contributed significantly to the functionality of this project. Their documentation and support were essential in implementing the gene loss database. We dedicate this work to the memory of **Bernardo Pinto**, whose enthusiasm and contributions during the early stages of this project were deeply valued. His joy and curiosity remain an inspiration to us.

