## Supplementary Table 1 for "Gene Loss DB: A curated database for gene loss in vertebrate species"

**Supplementary Table 1: Curated publications**

| Title | Nr. of GL | DOI |
|---|---|---|
| Functional or Vestigial? The Genomics of the Pineal Gland in Xenarthra. | 98 | 10.1007/s00239-021-10025-1 |
| Parallel Independent Losses of G-Type Lysozyme Genes in Hairless Aquatic Mammals. | 98 | 10.1093/gbe/evab201 |
| Recurrent loss of HMGCS2 shows that ketogenesis is not essential for the evolution of large mammalian brains. | 23 | 10.7554/eLife.38906 |
| Gene duplications and gene loss in the epidermal differentiation complex during the evolutionary land-to-water transition of cetaceans. | 79 | 10.1038/s41598-021-91863-3 |
| Analysis of the FGF gene family provides insights into aquatic adaptation in cetaceans. | 9 | 10.1038/srep40233 |
| Complete Inactivation of Sebum-Producing Genes Parallels the Loss of Sebaceous Glands in Cetacea. | 67 | 10.1093/molbev/msz068 |
| Genomewide analysis of sperm whale E2 ubiquitin conjugating enzyme genes. | 4 | 10.1007/s12041-021-01333-y |
| Molecular decay of the tooth gene Enamelin (ENAM) mirrors the loss of enamel in the fossil record of placental mammals. | 20 | 10.1371/journal.pgen.1000634 |
| Comparative genomics of sirenians reveals evolution of filaggrin and caspase-14 upon adaptation of the epidermis to aquatic life. | 10 | 10.1038/s41598-024-60099-2 |
| Epidermal cornification is preceded by the expression of a keratinocyte-specific set of pyroptosis-related genes. | 62 | 10.1038/s41598-017-17782-4 |
| Genetic basis of brain size evolution in cetaceans: insights from adaptive evolution of seven primary microcephaly (MCPH) genes. | 7 | 10.1186/s12862-017-1051-7 |
| Genetic evidence for the ancestral loss of short-wavelength-sensitive cone pigments in mysticete and odontocete cetaceans. | 16 | 10.1098/rspb.2002.2278 |
| Ancient convergent losses of yield potential risks for modern marine mammals. | 10 | 10.1126/science.aap7714 |
| A genomics approach reveals insights into the importance of gene losses for mammalian adaptations. | 111 | 10.1038/s41467-018-03667-1 |
| Positive Selection and Inactivation in the Vision and Hearing Genes of Cetaceans. | 42 | 10.1093/molbev/msaa070 |
| Progressive erosion of the Relaxin1 gene in bovids. | 43 | 10.1016/j.ygcen.2017.07.011 |
| Comparative genomics provides insights into the aquatic adaptations of mammals. | 25 | 10.1073/pnas.2106080118 |
| Inactivation of thermogenic UCP1 as a historical contingency in multiple placental mammal clades. | 31 | 10.1126/sciadv.1602878 |
| Convergent Loss of the Necroptosis Pathway in Disparate Mammalian Lineages Shapes Viruses Countermeasures. | 65 | 10.3389/fimmu.2021.747737 |
| Morphological and molecular evidence for a stepwise evolutionary transition from teeth to baleen in mysticete whales. | 11 | 10.1080/10635150701884632 |
| Unusual loss of chymosin in mammalian lineages parallels neo-natal immune transfer strategies. | 30 | 10.1016/j.ympev.2017.08.014 |

| | | |
|---|---|---|
| Patterns and tempo of PCSK9 pseudogenizations suggest an ancient divergence in mammalian cholesterol homeostasis mechanisms. | 165 | 10.1007/s10709-021-00113-x |
| The Novel Evolution of the Sperm Whale Genome. | 19 | 10.1093/gbe/evx187 |
| Convergent inactivation of the skin-specific C-C motif chemokine ligand 27 in mammalian evolution. | 18 | 10.1007/s00251-019-01114-z |
| Evolution of bitter taste receptors in humans and apes. | 47 | 10.1093/molbev/msi027 |
| Evolution of the MC5R gene in placental mammals with evidence for its inactivation in multiple lineages that lack sebaceous glands. | 17 | 10.1016/j.ympev.2017.12.010 |
| The dopamine receptor D gene shows signs of independent erosion in toothed and baleen whales. | 14 | 10.7717/peerj.7758 |
| The Singularity of Cetacea Behavior Parallels the Complete Inactivation of Melatonin Gene Modules. | 49 | 10.3390/genes10020121 |
| Genes lost during the transition from land to water in cetaceans highlight genomic changes associated with aquatic adaptations. | 266 | 10.1126/sciadv.aaw6671 |
| Increased rate of hair keratin gene loss in the cetacean lineage. | 28 | 10.1186/1471-2164-15-869 |
| Convergent Losses of TLR5 Suggest Altered Extracellular Flagellin Detection in Four Mammalian Lineages. | 23 | 10.1093/molbev/msaa058 |
| Regressed but Not Gone: Patterns of Vision Gene Loss and Retention in Subterranean Mammals. | 360 | 10.1093/icb/icy004 |
| Inactivation of the olfactory marker protein (OMP) gene in river dolphins and other odontocete cetaceans. | 12 | 10.1016/j.ympev.2017.01.020 |
| Cetacea are natural knockouts for IL20. | 11 | 10.1007/s00251-018-1071-5 |
| A drastic shift in the energetic landscape of toothed whale sperm cells. | 150 | 10.1016/j.cub.2021.05.062 |
| Molecular evolutionary analyses of tooth genes support sequential loss of enamel and teeth in baleen whales (Mysticeti). | 130 | 10.1016/j.ympev.2022.107463 |
| Phylogenetic profiling and gene expression studies implicate a primary role of PSORS1C2 in terminal differentiation of keratinocytes. | 11 | 10.1111/exd.13272 |
| Insights into the evolution of longevity from the bowhead whale genome. | 16 | 10.1016/j.celrep.2014.12.008 |
| Whale Hageman factor (factor XII): prevented production due to pseudogene conversion. | 1 | 10.1016/s0049-3848(97)00307-1 |
| GBA3: a polymorphic pseudogene in humans that experienced repeated gene loss during mammalian evolution. | 35 | 10.1038/s41598-020-68106-y |
| Aquatic adaptation and the evolution of smell and taste in whales. | 61 | 10.1186/s40851-014-0002-z |
| Pseudogenization of the tooth gene enamelysin (MMP20) in the common ancestor of extant baleen whales. | 9 | 10.1098/rspb.2010.1280 |
| Major taste loss in carnivorous mammals. | 12 | 10.1073/pnas.1118360109 |
| Birth-and-death evolution of ribonuclease 9 genes in Cetartiodactyla. | 59 | 10.1007/s11427-022-2195-x |
| Rod monochromacy and the coevolution of cetacean retinal opsins. | 34 | 10.1371/journal.pgen.1003432 |
| The loss of taste genes in cetaceans. | 75 | 10.1186/s12862-014-0218-8 |
| Mx1 and Mx2 key antiviral proteins are surprisingly lost in toothed whales. | 8 | 10.1073/pnas.1501844112 |
| Losses of human disease-associated genes in placental mammals. | 100 | 10.1093/nargab/lqz012 |
| Genomic and anatomical comparisons of skin support independent adaptation to life in water by cetaceans and hippos. | 230 | 10.1016/j.cub.2021.02.057 |
| Transition to an Aquatic Habitat Permitted the Repeated Loss of the Pleiotropic KLK8 Gene in Mammals. | 9 | 10.1093/gbe/evx239 |
| Loss or major reduction of umami taste sensation in pinnipeds. | 7 | 10.1007/s00114-012-0939-8 |

| | | |
|---|---|---|
| Decay of Skin-Specific Gene Modules in Pangolins. | 50 | 10.1007/s00239-023-10118-z |
| Convergent Cortistatin losses parallel modifications in circadian rhythmicity and energy homeostasis in Cetacea and other mammalian lineages. | 63 | 10.1016/j.ygeno.2020.11.002 |
| Rubbing Salt in the Wound: Molecular Evolutionary Analysis of Pain-Related Genes Reveals the Pain Adaptation of Cetaceans in Seawater. | 9 | 10.3390/ani12243571 |
| Comparative genomics analyses of alpha-keratins reveal insights into evolutionary adaptation of marine mammals. | 210 | 10.1186/s12983-017-0225-x |
