## Supplementary Table 2 for "Gene Loss DB: A curated database for gene loss in vertebrate species"

**Supplementary Table 2: Number of gloss annotations in Cetacea by species**

| Odontoceti | GLoss | Mysticeti | GLoss |
|---|---|---|---|
| *Tursiops truncatus* | 268 | *Balaenoptera acutorostrata* | 266 |
| *Physeter catodon* | 242 | *Balaena mysticetus* | 82 |
| *Orcinus orca* | 215 | *Balaenoptera bonaerensis* | 56 |
| *Lipotes vexillifer* | 98 | *Eschrichtius robustus* | 39 |
| *Neophocaena asiaeorientalis* | 63 | *Balaenoptera musculus* | 36 |
| *Delphinapterus leucas* | 42 | *Balaenoptera physalus* | 21 |
| *Phocoena sinus* | 38 | *Megaptera novaeangliae* | 20 |
| *Sousa chinensis* | 30 | *Caperea marginata* | 16 |
| *Kogia sima* | 24 | *Balaenoptera borealis* | 15 |
| *Lagenorhynchus obliquidens* | 24 | *Eubalaena japonica* | 15 |
| *Kogia breviceps* | 22 | *Eubalaena australis* | 13 |
| *Monodon monoceros* | 23 | *Eubalaena glacialis* | 13 |
| *Globicephala melas* | 21 | *Balaenoptera edeni* | 12 |
| *Phocoena phocoena* | 17 | *Balaenoptera brydei* | 1 |
| *Neophocaena phocaenoides* | 12 | *Balaenoptera omurai* | 1 |
| *Pontoporia blainvillei* | 14 | | |
| *Inia geoffrensis* | 14 | | |
| *Tursiops aduncus* | 14 | | |
| *Mesoplodon bidens* | 13 | | |
| *Ziphius cavirostris* | 12 | | |
| *Delphinus capensis* | 9 | | |
| *Peponocephala electra* | 8 | | |
| *Phocoenoides dalli* | 3 | | |
| *Platanista gangetica* | 3 | | |
| *Mesoplodon europaeus* | 3 | | |
| *Platanista minor* | 5 | | |
| *Mesoplodon densirostris* | 3 | | |
| *Mesoplodon mirus* | 2 | | |
| *Berardius bairdii* | 2 | | |
| *Mesoplodon ginkgodens* | 2 | | |
| *Mesoplodon carlhubbsi* | 2 | | |
| *Mesoplodon stejnegeri* | 2 | | |
| *Grampus griseus* | 2 | | |
| *Tasmacetus shepherdi* | 1 | | |
| *Pseudorca crassidens* | 1 | | |
| *Mesoplodon perrini* | 1 | | |
| *Globicephala macrorhynchus* | 1 | | |
| *Lagenorhynchus australis* | 1 | | |
| *Mesoplodon bowdoini* | 1 | | |
| *Hyperoodon ampullatus* | 1 | | |
| *Delphinus delphis* | 1 | | |
